# Compartment-specific opioid receptor signaling is selectively modulated by Dynorphin subtypes

**DOI:** 10.1101/2020.06.21.162206

**Authors:** Jennifer M. Kunselman, Achla Gupta, Ivone Gomes, Lakshmi A. Devi, Manoj A. Puthenveedu

## Abstract

Many signal transduction systems have an apparent redundancy built into them, where multiple physiological agonists activate the same receptors. Whether this is true redundancy, or whether this provides as-yet unrecognized specificity in downstream signaling, is not well understood. We address this question using the kappa opioid receptor (KOR), a physiologically relevant G protein-coupled receptor (GPCR) that is activated by multiple members of the Dynorphin family of opioid peptides. We show that, although highly related Dynorphins bind and activate KOR to similar extents on the cell surface, they localize KOR to distinct subcellular compartments, dictate different post-endocytic fates of the receptor, and differentially induce KOR signaling from the degradative pathway. Our results show that seemingly redundant endogenous opioid peptides that are often co-released can in fact fine-tune signaling by differentially regulating the subcellular spatial profile of GPCR localization and signaling.

## Introduction

The endogenous opioid system provides an excellent and physiologically relevant example to study redundancy in signaling systems in our body. Over twenty endogenous opioids have been identified, all of which preferentially activate one of three opioid receptors - delta, kappa, and mu opioid receptors - which are all members of the G protein-coupled receptor family of proteins (*1–3*). All these opioid peptides activate their cognate GPCRs broadly at similar levels in most of the readouts that have been measured (*4*). Whether all these opioid peptides are truly redundant or whether they contribute to signaling diversity in some unknown manner, has been a long-standing question in the field.

Beyond initial signaling from the cell surface, subsequent phases of signaling from intracellular compartments is emerging as a key determinant of the downstream consequences of receptor activation (*5–7*). While this is still an emerging field, a growing body of evidence suggests that GPCRs are active in endosomes and other intracellular compartments, and that receptors in endosomes can cause distinct signaling consequences compared to receptors on the cell surface (*8–10*). Receptors rapidly and dynamically move between intracellular compartments and the surface by trafficking, and trafficking could therefore act as a master regulator of GPCR signaling by selectively amplifying signals from specific locations (*11–12*). Whether physiological systems take advantage of trafficking to localize receptors to different compartments and dictate location-biased signaling, however, is still not known. Here we use the kappa opioid receptor (KOR) as a model GPCR to whether Dynorphin peptides - seemingly redundant physiological agonists - could in fact drive spatially restricted signaling outcomes based on the particular Dynorphin that activates KOR.

## Results

We first focused on four physiologically relevant endogenous Dynorphin peptides - Dynorphin A17 (DynA), Dynorphin A8 (DynA8), Dynorphin B (DynB), and α-neoendorphin (α-neo). These peptides differ mainly in the length and sequence of the peptide at the C-termini (Figure 1A). Consistent with previous findings, these peptides bound KOR at relatively comparable affinities (Table 1). Additionally, the Dynorphin peptides inhibited whole-cell cAMP levels to comparable levels (Figure 1B). Further, the rates and extents of agonist-mediated KOR clustering into endocytic domains, as well as receptor endocytosis, when measured in neuroendocrine cells stably expressing KOR tagged with a pH-sensitive GFP (*13*) (SpH-KOR), were similar between these peptide agonists (Figure 1C-D). These results indicate that different Dynorphin peptides activate and endocytose KOR to similar levels, consistent with previous findings (*14*).

**Figure 1.**
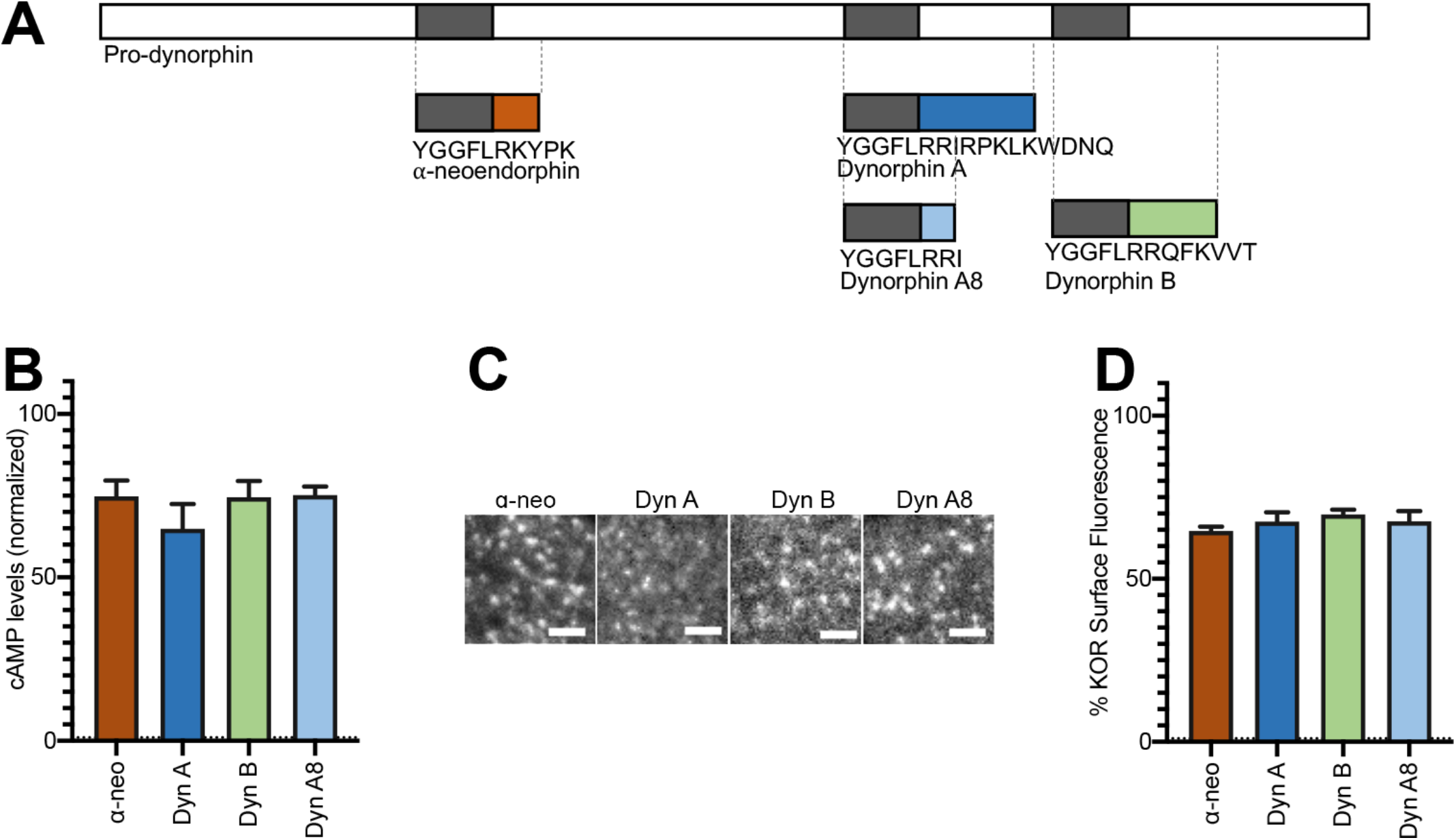
Initial activation and internalization of KOR by Dynorphins are comparable. **A)** Schematic of the regions of Pro-dynorphin from which four key peptides are generated, showing that DynA and DynB are processed from immediately adjacent regions. **B)** Initial KOR signaling, measured as whole-cell cAMP inhibition compared to control (Ctrl), is similar across Dynorphins in PC12 cells stably expressing SpH-KOR. **C)** Representative images of PC12 cells stably expressing SpH-KOR, imaged in TIR-FM, show that agonist-mediated receptor clustering on the cell surface, 1 min after treatment, is similar between all four peptides. Scale bar = 2μm. **D)** Quantitation of the loss of surface SpH-KOR fluorescence, as an index of internalization, after 5 min of treatment with each peptide, normalized to surface fluorescence before agonist, show similar levels of internalization for all four peptides. DynA: n=10 cells; DynB: n=10 cells; DynA8: n=10 cells; α-neo: n=11 cells.

**Table 1.** Displacement binding parameters for Dynorphin peptides at PC12SPH-kOR and CHO-kOR cells. Displacement binding assays were carried out with [^3^H] diprenorphine (3 nM) using membranes from PC12SPH-kOR (100 mg) and CHO-kOR (15 mg) cells without or with different concentrations (10^-12^ - 10^-5^ M) of Dyn A8, Dyn A17, Dyn B13 or a-neo-endorphin in 50 mM Tris-Cl buffer containing 100 mM NaCl, 10 mM MgCl2, 0.2 mM EGTA and protease inhibitor cocktail as described in Methods. Data are mean ± SE (n=3). *p<0.05, **p<0.01, ***p<0.001,****p<0.0001 v/s pH 7.4, One-Way ANOVA. n_H_, % receptors in high affinity state.

| Ligand |  | PC12SPH-κOR cells |  |  |  | CHO-κOR cells |  |  |  |
| --- | --- | --- | --- | --- | --- | --- | --- | --- | --- |
|  |  | Low K<sub>i</sub> (nM) | High K<sub>i</sub> (nM) | % bound at 10 μM | n<sub>H</sub> | Low K<sub>i</sub> (nM) | High K<sub>i</sub> (nM) | % bound at 10 μM | n<sub>H</sub> |
| Dyn A8 | pH 7.4 | 56.9±0.3 | 0.020±1.90 | 18.30±1.48 | 36.9% | 563±0.1 | 0.41±0.15 | 21.51±1.54 | 41.6% |
|  | pH 6.5 | 62.9±0.2**** | 0.024±0.35 | 22.84±1.54 | 39.6% | 634±0.2**** | 0.54±0.19 | 22.61±1.64 | 43.8% |
|  | pH 5.5 | 92.8±0.2**** | 0.040±0.34 | 27.11±1.92** | 29.6% | 778±0.2**** | 0.89±0.23 | 27.09±1.29* | 28.5% |
| Dyn A17 | pH 7.4 | 52.4±0.3 | 0.016±0.47 | 15.22±1.56 | 39.1% | 119±0.2 | 0.26±0.54 | 18.72±1.09 | 29.4% |
|  | pH 6.5 | 63.6±0.2**** | 0.020±0.40 | 20.72±1.64* | 41.2% | 147±0.2**** | 0.34±0.49 | 19.13±0.67 | 31.1% |
|  | pH 5.5 | 65.9±0.2**** | 0.032±0.57 | 23.12±1.45** | 39.4% | 194±0.1**** | 0.57±0.22 | 23.39±1.57* | 33.5% |
| Dyn B13 | pH 7.4 | 37.8±0.2 | 0.010±0.31 | 11.13±1.07 | 33.4% | 355±0.1 | 0.38±0.11 | 17.16±0.82 | 42.2% |
|  | pH 6.5 | 40.5±0.2**** | 0.011±0.39 | 18.52±1.32*** | 42.4% | 382±0.2**** | 0.49±0.24 | 21.12±1.04* | 42.7% |
|  | pH 5.5 | 48.5±0.2**** | 0.014±0.44 | 24.96±1.14**** | 40.9% | 406±0.1**** | 0.60±0.14 | 23.73±1.25*** | 42.7% |
| α-neo-endorphin | pH 7.4 | 39.1±0.1 | 0.011±0.20 | 13.32±1.84 | 38.7% | 591±0.1 | 0.18±0.34 | 20.55±1.57 | 21.4% |
|  | pH 6.5 | 50.2±0.2**** | 0.013±0.33 | 19.69±1.08* | 45.2% | 693±0.2**** | 0.29±0.39 | 21.02±1.45 | 27.9% |
|  | pH 5.5 | 77.1±0.2**** | 0.016±0.31 | 29.42±1.81**** | 40.8% | 867±0.1**** | 0.35±0.12 | 25.72±0.83* | 23.5% |

We next asked if different Dynorphins selectively regulate the fate of KOR after initial activation. We first tested post-endocytic recycling of KOR, by adapting a discrete imaging method to quantitate the rate of individual KOR recycling events over unit time. SpH-KOR fluorescence is quenched in acidic endosomal compartments and is rapidly dequenched when receptors recycle back to the cell surface and are exposed to the extracellular media. This dequenching can be visualized at single-event levels using Total Internal Reflection Fluorescence microscopy (TIR-FM), where these recycling events show a distinctive sudden spike in fluorescence followed by an exponential decay as the receptors diffuse on the cell membrane (Figure 2A-C). This method allows us to quantitate individual recycling events in the same cells over time without the confounding effects of continuing endocytosis (*15*). When the number of SpH-KOR recycling events were quantitated, normalized to time and cell area, a significantly higher number of recycling events was seen after DynB-induced KOR internalization, compared to DynA, DynA8, or α-neo (Figure 2D).

**Figure 2.**
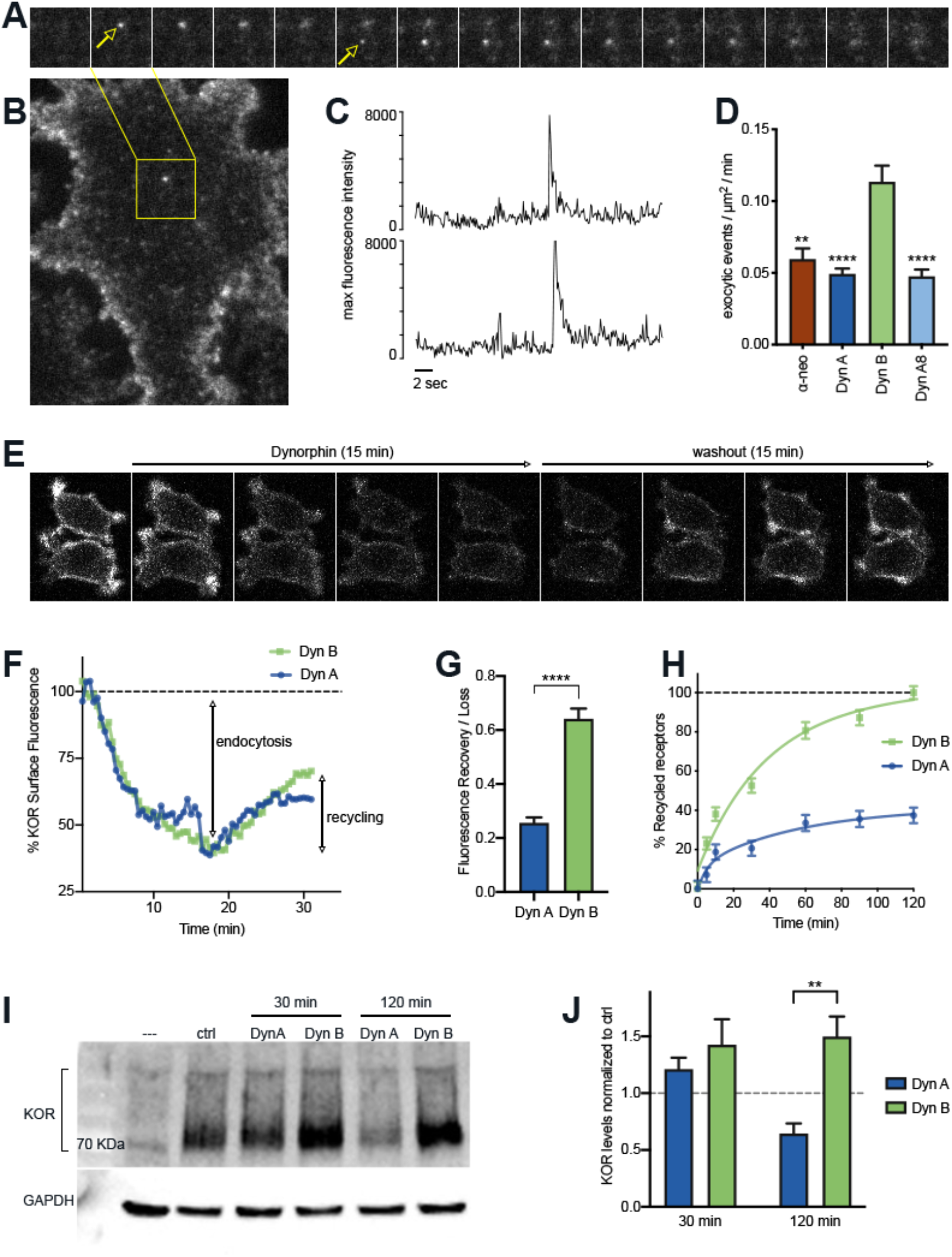
The post-endocytic fate of KOR is determined by the specific Dynorphin that activates it. **A)** Frames from a time lapse movie of a representative region of a PC12 cell stably expressing SpH-KOR shown in **(B),** treated for 5 min with DynB and imaged in TIR-FM, showing two example exocytic events mediating KOR recycling. **C)** Fluorescence traces of the two exocytic events showing a characteristic abrupt increase in maximum fluorescence intensity followed by exponential decay. **D)** Quantitation of the number of exocytic events per μm^2^ per min showing a significant increase in the number for DynB compared to the other peptides (**p<.01, ****p<.0001 in multiple comparisons after ANOVA, n= 14, 39, 52, and 33 respectively). **E)** Ensemble SpH-KOR surface fluorescence, measured over time using confocal microscopy, decreases upon Dynorphin addition because of quenching of internalized SpH-KOR, and an increase upon Dynorphin washout as receptors recycle to the surface and SpH-KOR is dequenched. **F)** Quantification of change in ensemble surface fluorescence over 30 minutes following Dynorphin A or B treatment, normalized to fluorescence before agonist addition, showing the loss during endocytosis and increase during recycling. **G)** Quantitation of the amount of SpH-KOR recycled, normalized to the amount endocytosed, after DynA or DynB, show that a higher amount of receptor recycled after DynB. (****p<0.0001 by Mann-Whitney; n=33 and 30 fields for DynA and DynB respectively). **H)** Recycling of KOR to the cell surface after DynA or DynB measured by ELISA show a much higher rate and extent of recycling after DynB. **I)** Representative immunoblot of total receptor levels in SpH-KOR PC12 cells, treated with cycloheximide for 2 hours prior to DynA or DynB treatment for the indicated times, show loss of receptor after 120 min of DynA but not DynB treatment. GAPDH is shown as a control. **J)** Quantification of total receptor levels normalized to untreated control cells under each condition (**p<.01 by post-hoc comparison after two-way ANOVA; n=5).

To test whether this increase in the rate of discrete recycling events corresponded to an increase in receptor levels at the cell surface, we measured ensemble changes in surface KOR levels by two different methods - whole-cell fluorescence and ELISA-based methods. We focused on Dynorphin B and Dynorphin A as a highly relevant and interesting pair, as both are processed from immediately adjacent regions of pro-dynorphin and are often co-expressed in physiologically relevant brain regions (*16–17*). When SpH-KOR fluorescence was followed live by confocal imaging, surface fluorescence decreased after Dynorphin addition, as was expected with internalization. The fluorescence decrease reached a plateau at 10 minutes, suggesting that an equilibrium between endocytosis and recycling was reached at that time (Figure 2E). When agonist was removed by washing out the media, to specifically measure recycling without the contribution of endocytosis, the fluorescence rates recovered to higher levels in cells activated with DynB than with Dyn A (Figure 2F-G). These results were supported by ELISA-based assays that measured surface KOR levels. After Dynorphin washout, DynB-treated cells showed a higher rate of KOR recovery on the cell surface compared to DynA (Figure 2H). These ensemble measurements support the single-event analysis in that DynB causes KOR to recycle at a higher rate than DynA.

Because KOR did not recycle efficiently when activated by DynA, we next asked whether DynA-activated KOR was sorted into the degradative pathway. To test this, PC12 cells expressing SpH-KOR were pretreated with cycloheximide 2 hours before agonist addition to inhibit any new protein synthesis and to measure agonist-mediated turnover of KOR. Total KOR levels were determined through immunoblotting, after DynA or DynB treatment (Figure 2I). When total receptor levels were quantified, DynA treatment caused a loss of 50% of KOR 2 hours after treatment, while DynB treatment caused no loss at the same time point (Figure 2J). These results suggest that, after endocytosis, DynA preferentially sorts KOR into the degradative pathway, while DynB sorts KOR into the recycling pathway.

Considering the emerging importance of spatial encoding in diversifying the outcomes of GPCR signaling (*7, 11–12*), we next asked whether DynA or DynB generated distinctive intracellular localization patterns of KOR. PC12 cells stably expressing SpH-KOR were treated with DynA or DynB for 20 min. They were then fixed and stained for APPL1 (very early endosomes), EEA1/Rab5 (early sorting endosomes), Rab11 (recycling endosomes), Rab7 (late endosomes), and Lamp1 (lysosomes), as markers for the biochemically distinct compartments along the early, recycling, and late endosomal pathway. KOR colocalized predominantly with Rab7, a component of the late endocytic pathway, when activated by DynA (example in Figure 3A). In contrast, KOR colocalized predominantly with Rab11, a component of the recycling endocytic pathway, when activated by DynB. Using automated object-picking, we quantitated the fraction of KOR endosomes that colocalized with each endosomal marker in these conditions. KOR localized mainly to compartments marked by Rab7 and Lamp1 when activated by DynA, but to compartments marked by EEA1, Rab5, and Rab11 when activated by DynB (Figure 3B-C). These results show that KOR is concentrated in different endosomal compartments based on the Dynorphin that activates it.

**Figure 3.**
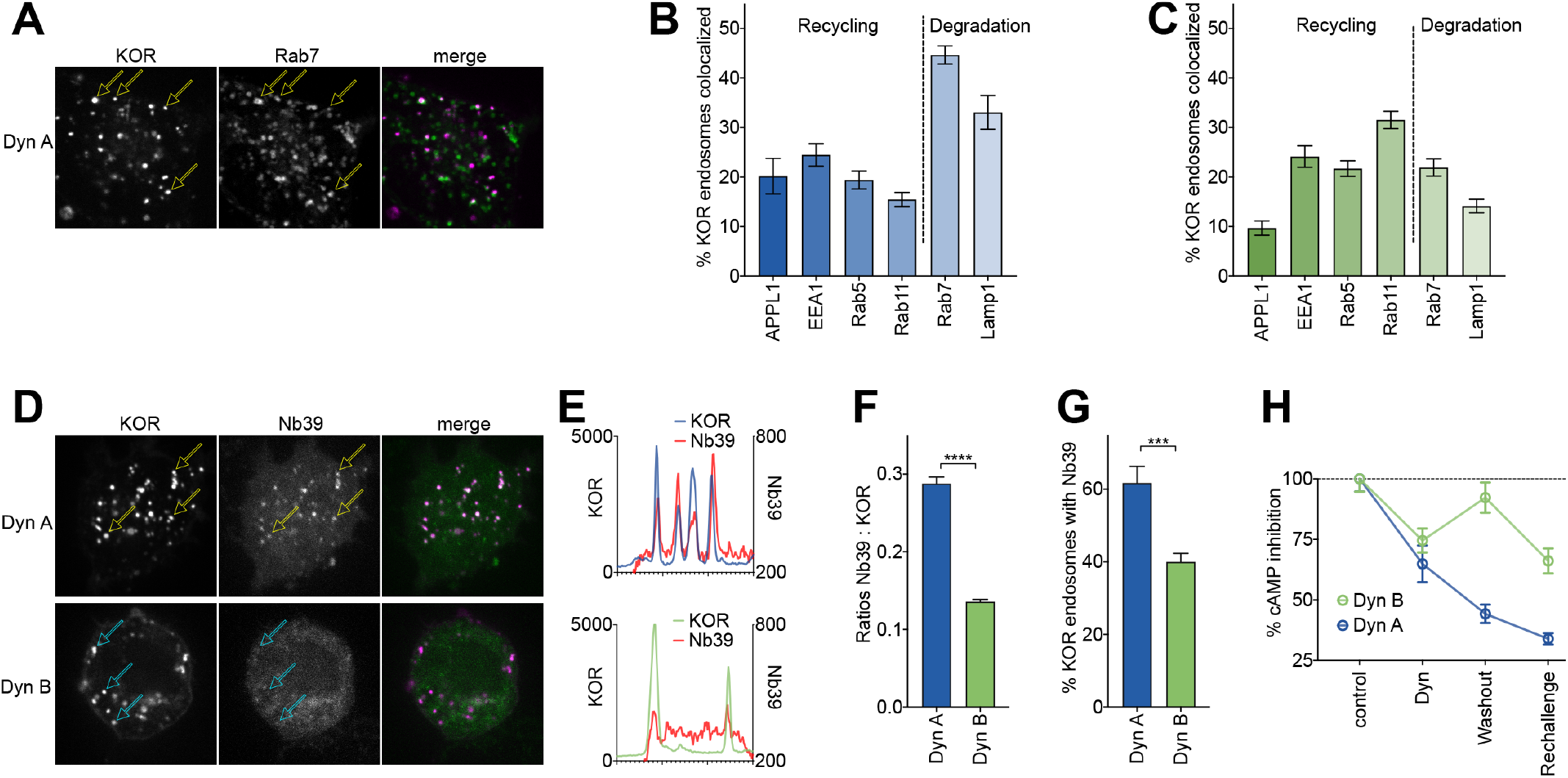
DynA selectively drives KOR signaling from late endosomal compartments. **A)** Representative image of a PC12 cell expressing FLAG-KOR and Rab7-GFP, treated with DynA for 20 minutes. Yellow arrows denote KOR endosomes that colocalize with Rab7. **B)** SpH-KOR cells treated with DynA for 20 min were fixed and processed for immunofluorescence with the noted markers. Quantitation, across multiple cells, of the percentage of KOR endosomes that colocalize with each the endosomal markers noted. KOR primarily localizes in Rab7 and Lamp1 positive late endosomes after DynA (n= 8, 10, 9, 11, 20, and 17 cells respectively). **C)** A similar quantitation of immunofluorescence images after DynB shows that KOR localizes less with late endosomes, and more with markers of early/recycling endosomes (n= 18, 16, 15, 18, 23, and 29 cells respectively). **D)** Representative images of PC12 cells expressing FLAG-KOR and Nb39, imaged live after treatment with DynA or DynB for 20 min. Yellow arrows in DynA show KOR endosomes that recruited Nb39, while cyan arrows in DynB show KOR endosomes that do not show obvious concentration of Nb39. **E)** Linear profile plots of fluorescence of KOR and Nb39, measured along lines drawn across regions of the cell with KOR endosomes after DynA or DynB treatment for 20 min, show that Nb39 fluorescence increases along with KOR in DynA, but less noticeably with DynB. **F)** Ratios of integrated fluorescence of Nb39:KOR in endosomes identified by 3D object analysis show higher amounts of Nb39 relative to KOR in DynA-treated cells (****p<.0001 by Mann Whitney, n=766 and 800 endosomes for DynA and DynB respectively). **G)** Quantitation of the percentage of KOR endosomes per cell with a noticeable increase in Nb39 fluorescence above background, shows a higher fraction of KOR endosomes recruited Nb39 in DynA-treated cells (***p<.001 by Mann Whitney, n= 11 and 14 cells for DynA and DynB respectively). **H)** cAMP levels after initial Dynorphin treatment for 5 min, washout for 25 min, or a Dynorphin rechallenge at end of the washout, show comparable initial cAMP inhibition by both DynA and DynB, but persistent signaling by DynA after agonist washout.

The agonist-selective localization of KOR to specific endosomes raised the exciting possibility that different Dynorphins could generate distinct subcellular spatial patterns of KOR signaling. To test this possibility, we combined conformation-selective biosensors, which are revolutionizing the field, and high-resolution imaging to ask whether KOR was active in endosomes. A nanobody (Nb39) that specifically recognizes the active conformation of KOR (*18*), when co-expressed with FLAG-tagged KOR in PC12 cells, localized efficiently to endosomes that also contained DynA-activated KOR. In contrast, Nb39 localized less to endosomes containing DynB-activated KOR (Figure 3D-E). When the fraction of total number of KOR endosomes per cell that colocalized with Nb39 was quantitated by analyzing 3D stacks, endosomes containing DynA activated KOR recruited Nb39 at a significantly higher level (Figure 3F-G), suggesting that KOR was in an active conformation in the endosomes specifically after activation by DynA. Consistent with this idea, DynA, but not DynB, retained cAMP inhibition even after surface receptors were removed by agonist washout (Figure 3H). Together, our results suggest that DynA, but not DynB, specifically coordinates activation and cAMP inhibition by KOR on late endosomes and lysosomes.

Importantly, this Dynorphin-selective coordination of KOR recycling and endosomal activation was conserved in striatal neurons. To directly measure KOR recycling, E18 rat primary embryonic striatal neurons were transfected with SpH-KOR, and individual recycling events were imaged in TIRFM. The number of individual exocytic events, when quantified per minute and normalized to cell area, was significantly lower in neurons treated with DynA compared to DynB (Figure 4A). This suggested that DynB, but not DynA, preferentially sorted KOR to recycling endosomes in neurons. We directly tested this by detecting the steady-state localization of KOR in endosomes after DynA or DynB treatment. KOR colocalized predominantly with Rab7 when activated by DynA, and with Rab11 when activated by DynB (Figure 4B-C). To test whether this differential localization correlated with differential location-based activation of KOR in endosomes, we expressed Nb33, a distinct nanobody that also recognizes that active conformation of opioid receptors (*19*), fused to GFP, in neurons. Endosomes containing DynA-activated KOR recruited Nb33, while endosomes containing DynB-activated KOR recruited Nb33 to a noticeably lesser extent. This recruitment was readily apparent in dendritic projections, where endosomes were distinctly visible (Figure 4D). The percentage of KOR endosomes that recruited Nb33 was significantly higher for DynA-activated KOR than for DynB (Figure 4E), showing that dynorphin-selective spatial activation of KOR was conserved in neurons.

**Figure 4.**
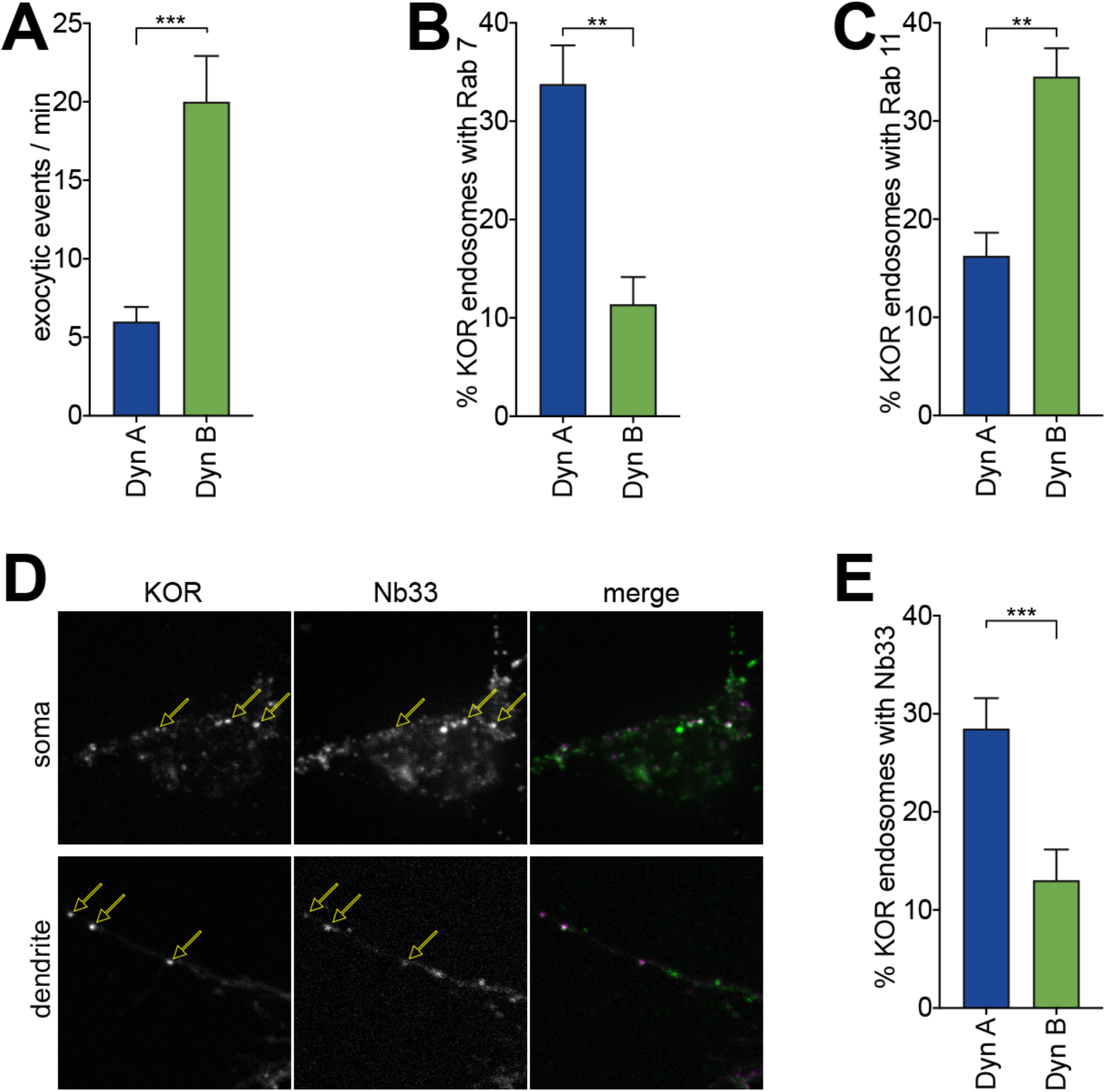
DynA-specific late endosomal localization and signaling is conserved in striatal neurons. **A)** The number of discrete exocytic events quantitated in rat medium spiny neuron (MSN) expressing SpH-KOR, show increased recycling for DynB compared to the other peptides (***p<.001, n=8 cells). **B)** Quantification of the percentage of KOR endosomes colocalized with of Rab7 in MSN expressing SpH-KOR treated with DynA or DynB (**p<.01, n=5 cells). **C)** Quantification of the percentage of KOR endosomes colocalized with of Rab11 in MSN expressing SpH-KOR treated with DynA or DynB (**p<.01, n=5 and 9 cells). **D)** Colocalization of FLAG-KOR and Nb33-GFP in the soma and in dendrites of MSNs treated with Dyn A, seen by confocal microscopy. Yellow arrows show KOR endosomes that recruit Nb33. **E)** Quantitation of the percentage of KOR endosomes per cell with a noticeable increase in Nb33 fluorescence above background, shows a higher fraction of KOR endosomes recruited Nb33 in DynA-treated cells (**p=0<.001, n=10 cells). All p values were from non-parametric Mann Whitney tests.

## Discussion

Together, our results reveal an unanticipated difference between physiologically important endogenous opioid peptides in encoding the subcellular spatial patterns of KOR signaling. The exact mechanism by which KOR is localized to different compartments is not clear, although it is likely mediated by specific interactions of the unstructured cytoplasmic tail with trafficking proteins like the PDZ-interacting protein NHERF1/EBP50 (*20*). In this context, different Dynorphin peptides could lock KOR into conformations that selectively allow or inhibit interactions with signaling proteins, essentially defining the receptor interactome in an agonist-specific manner. This conformational lock could require the presence of the ligand that is co-internalized with the receptor, although an exciting possibility is that the ligands provide a “conformational memory” to KOR that is sustained through the endocytic trafficking pathway. At present it is not clear what could provide such a conformational memory. It is possible that different Dynorphins could cause agonist-specific post-translational modifications, which is a general emerging theme for opioid receptors (*21–22*). To the best of our knowledge, however, this is the first example of different physiological agonists driving spatial localization and trafficking of a GPCR - a process that has been studied largely using receptor mutants or by depleting key components of the trafficking machinery so far (*9, 11, 23*).

It is surprising that DynA, which drives receptors into the late endosomal and degradative pathway, caused Nb33/Nb39 recruitment to endosomes. For most other canonical GPCRs studied, early endosomes are the main compartments that mediate endosomal signaling. Nevertheless, the idea that KOR in the late endosomal pathway activates Gi to reduce cAMP is consistent with the presence and activation of Gi on lysosomes by other GPCRs, as shown for cannabinoid receptors trafficking from the Golgi (*24*). DynA activated KOR could also activate alternate pathways, such as mTOR signaling, on late endosomes and lysosomes. Interestingly, mTOR signaling is involved in several functional effects of KOR (*25–27*). Because mTOR-mediated effects of KOR are primarily deleterious, there have been efforts to generate chemical agonists that bypass this signaling pathway. It is possible that mTOR signaling downstream of KOR is location-specific to DynA. This provides key support to the emerging and exciting idea that location-based activation/inhibition could be a strategy for generating biased agonists that maximize therapeutic potential and minimize adverse effects.

Physiological systems could leverage receptor sorting to fine-tune both spatial and temporal aspects of GPCR signaling. In the case of KOR, because DynA drives very little KOR to recycle, but promotes endosomal KOR activation, the net effect would be to cause a sustained cAMP inhibition from endosomes after a single exposure. Because DynB drives rapid KOR recycling and promotes endosomal signaling to a lesser extent, the net effect would be to cause short-lived cAMP inhibition primarily from the surface. On the other hand, the rapid recycling and resensitization caused by DynB, unlike DynA, would sensitize cells to repeated pulses of ligand release. Whether these different Dynorphins are always co-released in the nervous system, or whether different brain regions selectively release specific Dynorphins, is still unclear. DynA and DynB are generated from prodynorphin likely in the late stages of dense core vesicle maturation and could be predominantly co-released, but, there could be mechanisms that actively segregate or selectively release Dynorphins. In any case, KOR is activated by many opioid peptides that are generated from multiple precursor peptides, with distinct consequences due to signaling from the cell surface (*28*). Our results, that highly related opioid peptides regulate spatial encoding of KOR, therefore suggest an unanticipated layer of granularity to the anatomical and functional maps of the brain.

## Acknowledgements

We thank Dr. Dan Shiwarski and Marlena Darr for essential initial technical assistance with the project. We thank Drs. Robert Fuller, Carole Parent, Alan Smrcka, and Lloyd Fricker for expert discussions. JMK was supported by NIH T-32-GM007315, LAD by NIH NS026880 and DA008863, and MAP by NIH GM117425 and by NSF 1935926.

## Materials and Methods

### Reagents, Constructs, and Cells

Dynorphin A, Dynorphin B, Dynorphin A (1-8), and α-neoendorphin (used at 1μM) were purchased from Tocris Bioscience and/or Phoenix Pharmaceuticals. Naltrexone (10μm) was purchased from Sigma Aldrich (St. Louis, MO). Stable non-clonal PC12 cells expressing superecliptic phluorin (SpH)-KOR were selected in puromycin (Gibco) and grown in F12K media supplemented with 10% horse serum and 5% fetal bovine serum (Gibco) in collagen coated flasks. PC12 cells were also transiently transfected with KOR fused to FLAG on its n terminus using Lipofectamine 2000 transfection kit (ThermoFisher). Transfected cells were imaged two-three days after transfection. E18 rat striatal neurons were obtained from BraintBits LLC and cultured on poly-d-lysine (Sigma) coated coverslips for 1 week in Neurobasal media (Gibco) supplemented with B27 (Gibco) and 1% Glutamax (Gibco) and 1% PenicillinStreptomycin (Gibco) before transfection.

### Live Cell Imaging

Cells were plated to poly-D-lysine (Sigma) coated 25mm coverslips. Cells were imaged two days later in Leibovitz L15 imaging medium (Gibco) and 1% fetal bovine serum at 37 °C in a CO_2_-controlled imaging chamber, using a Nikon Eclipse Ti automated inverted microscope with a 60x or a 100x 1.49 N.A. TIRF objective or a 20x 0.75 N.A. objective. Images were acquired with an iXon+ 897 electron-multiplying charge-coupled device camera with a solid state laser of 488 nm or 647 nm as a light source. Images were analyzed using FIJI (Schindelin et al., 2012).

### Quantification of Individual Recycling Events

PC12 cells stably expressing SpH-KOR were treated with KOR agonists: Dyn A17, Dyn B13, Dyn A8, or α-neo endorphin for 5 minutes to induce receptor clustering and internalization at 37°C. Receptor clustering was visualized using TIRF microscopy. Images were acquired every 3 seconds for a total of 5 minutes. Following internalization, a recycling movie was recorded at 10 Hz for 1 minute in TIRF. The number of exocytic recycling events were manually scored in FIJI to determine the recycling rate for each agonist. Recycling events were also recorded using the same method In primary striatal rat medium spiny neurons that were transfected with the SpH-KOR plasmid. Recycling movies were taken 30 minutes after agonist addition in neurons. Statistical significance was determined using a one-way ANOVA.

### Ensemble Recycling Assay

Receptor surface levels were measured in PC12 cells stably expressing SpH-KOR by using confocal microscopy on a 20x objective and 488-nm laser. Images were collected in 30 second intervals across 20 different cell fields. After 2 minutes of baseline an agonist (Dyn A17 or Dyn B13) was added to imaging media. Following agonist addition, images were collected for 15 minutes. After 15 minutes, agonist was removed and the imaging media was replaced with fresh media containing antagonist (naltrexone). Images were then collected for another 15 minutes. Fluorescence intensities were corrected by a background threshold and normalized by the average fluorescence of the baseline frames before agonist treatment. Surface fluorescence analysis was conducted using an ImageJ Macro automated script (National Institutes of Health) (Weinberg, 2019). Fluorescence recovery/loss ratios after washout were quantified by normalizing the fluorescence values after washout to the total fluorescence lost before washout. Cell fields that did not respond to Dynorphin treatment were excluded from analysis. Statistical significance was determined by using a student’s paired t test comparing the endpoints between agonist treatment.

### Immunofluorescence of Endosomal Markers

PC12 cells stably expressing SpH-KOR were plated on poly-d-lysine (Sigma Aldrich) coverslips and grown for 24-48 hours at 37 °C. Cells were then incubated with different agonists (Dyn A, Dyn B, Dyn A8, or α-neo for 20 minutes at 37 °C. Cells were then fixed with 4% paraformaldehyde (PFA), pH 7.4, for 20 minutes. Cells were then rinsed with complete PBS for a total of 2 times and then blocked in PBS containing calcium, magnesium, with 5% FBS, 5% 1M glycine, and 0.75% Triton X-100. SpH-KOR cells were then incubated with an antibody for one of the endosomal markers (anti-APPL1, -EEA1, -Rab5, - Rab11, -Rab7, -Lamp1 Rabbit mAB) (Cell Signaling Technology) for 1 hour. Cells were washed three times with PBS containing calcium and magnesium and then labeled with Alexa 647 goat anti-rabbit secondary antibody (1:1000) in a blocking buffer for 1 hour. Confocal imaging of cells was performed using spinning disk confocal microscope (Andor) and 100x objective. Representative images were taken across 10-20 fields for each agonist treatment and endosomal marker. Three biological replicates were performed in each condition.

### Endosomal Colocalization Quantification

The percent colocalization of the endosomal marker with the total number of receptor positive endosomes was determined using an ImageJ Macro: Object.picker (Weinberg, 2020) (doi.10.5281/zenodo.3811031) to identify the total number of endosomes containing receptor in one channel and determine the colocalization with an endosomal marker in another channel. The Image J macro: 3D Object Counter was used as another method of quantification for colocalization. Integrated density values for each object detected in both the receptor and endosome marker channels were used to determine a ratio of endosomal colocalization by dividing the endosomal marker signal by the receptor signal.

### Immunoblotting

PC12 cells stably expressing SpH-KOR were grown in a PDL coated 12-well plate for 2 days at 37 °C. Cells were treated with cycloheximide (3μg/ml) for 2 hr before agonist incubation. Cells were treated with Dyn A17 or Dyn B13 for 30 min or 2hr. A non-agonist treated well of PC12 cells stably expressing SpH-KOR and a well of PC12 cells not expressing SpH-KOR were used as controls. Following agonist treatments, cells were placed on ice and rinsed twice with PBS containing calcium and magnesium. Cells were directly lysed in the plate using 2X RSB (Bio-Rad, Hercules, CA). Lysates were placed on ice for 30 minutes and then sonicated in 5 second pulses. Following sonication, lysates were incubated at 37 °C for 1 hour. Lysates were run on 10% stain-free gels (BioRad), which were then transferred to nitrocellulose membrane overnight. Membranes were blocked in 5% milk and then probed with anti-GFP Chicken pAB (Abcam) to detect total receptor levels in each condition. Blots were developed using the iBright imager for chemiluminescence signal and quantified using FIJI software. Receptor signal for each condition was normalized to the no treatment control. Five biological replicates were performed. Statistical analysis was performed using two-way ANOVA across time and drug treatment.

